# GUIdEStaR (G-quadruplex, uORF, IRES, Epigenetics, Small RNA, Repeats), the integrated metadatabase in conjunction with neural network methods

**DOI:** 10.1101/2021.02.25.432957

**Authors:** Jee Eun Kang

## Abstract

GUIdEStaR integrates existing databases of various types of G-quadruplex, upstream Open Reading Frame (uORF), Internal Ribosome Entry Site (IRES), methylation to RNA and histone protein, small RNA, and repeats. GUIdEStaR consists of approx. 40,000 genes and 320,000 transcripts. An mRNA transcript is divided into 5 regions (5’UTR, 3’UTR, exon, intron, and biological region) where each region contains presence-absence data of 169 different types of elements. Recently, artificial intelligence (AI) based analysis of sequencing data has been gaining popularity in the area of bioinformatics. GUIdEStaR generates datasets that can be used as inputs to AI methods. At the GUIdEStaR homepage, users submit gene symbols by clicking a “Send” button, and shortly result files in CSV format are available for download at the result website. Users have an option to send the result files to their email addresses. Additionally, the entire database and the example Java codes are also freely available for download. Here, we demonstrate the database usage with three neural network classification studies-1) small RNA study for classifying transcription factor (TF) genes into either one of TF mediated by small RNA originated from SARS-CoV-2 or by human microRNA (miRNA), 2) cell membrane receptor study for classifying receptor genes as either with virus interaction or without one, and 3) nonsense mediated mRNA decay (NMD) study for classifying cell membrane and nuclear receptors as either NMD target or non-target. GUIdEStaR is available for access to the easy-to-use web-based database at www.guidestar.kr and for download at https://sourceforge.net/projects/guidestar.

## Introduction

Varied types of sequence compositions, structures, and epigenetic modifications are interwoven for orchestrating gene functions. Here, we introduce the integrative database, GUIdEStaR that unites existing databases of G-quadruplex (GQ) provided by Chariker et al., uORF from uORF-Tools, IRES from IRESbase, Epigenetic modification from Met-DB V2.0 and Choi et al., Small RNA from DASHR 2.0, and Repeats from RepeatMasker [1-7]. These elements are interdependent in controlling gene regulation through functional interactions. GQ, which is a topological change required of numerous conserved regulatory elements, has become increasingly important in recent years as the mutual dependency between GQ and epigenetic modification has been unveiled [8,9]. uORFs with a conserved peptide are often cleaved by miRNA for controlling the translation of their main ORFs that are frequently regulatory genes such as transcription factors and 5’ signal transduction factors [10-12]. They, together with IRES are known to select a specific RNA variant for translation and cellular localization [13]. Genes containing small RNA loci implicate various functions in cellular signaling, and the binding affinity of small RNA is regulated by epigenetic modification [14-16]. And repeats function as a gene regulator; for example, downregulation caused by intronic repeats can be suppressed by epigenetic modification [17,18]. Finally, epigenetic modification regulates gene expression, biogenesis of miRNA, and suppression of transposable elements [19,20].

Currently, there are no integrative bioinformatics resources that offer users with a means to study the associations of these elements. GUIdEStaR provides an effective analysis tool for studying interdependences of these elements. With one-click process at the GUIdEStaR homepage, users are given the access to the presence-absence data of 845 elements per transcript, which can be inputted to neural methods. GUIdEStaR consists of approx. 40,000 genes and 320,000 transcripts.

Here, we conducted three studies to demonstrate how to chain GUIdEStaR with neural network methods-classification and attribute selection. The first study was to classify human transcription factors (TFs) into either one of TF mediated by small RNA that has its origin in coronavirus or mediated by human miRNA [21, 22]. In addition to GUIdEStaR, the datasets included log base2 (fold change in gene level abundance) (log base2 FC) information in eight different cell types (ciliated cells, basal cells, club cells, goblet cells, neuroendocrine cells, ionocytes, and tuft cells) from single-cell (sc) RNA-seq [23,24]. The top ranked attributes included epigenetic modification to Histone H3, piRNA, short interspersed nuclear element (SINE) families, long interspersed nuclear element (LINE) families, epigenetic modification-RNA N6-methyl-adenosine (m6A) target sites of WTAP, METTL14, and METTL3, and log base2 FC information. The binary neural network models achieved the average accuracy of 90% (Table 1, Table S1, supplementary data).

**Table 1.** Accuracies of binary classifiers

| Accuracy | 5'UTR | Biol. Reg. | Exon | Intron | 3'UTR |
| --- | --- | --- | --- | --- | --- |
| Small RNA targets<br>(virus origin vs. human origin) | 88 % | 88 % | 93 % | 91 % | 89 % |
| Receptors with virus interaction vs.<br>receptors without virus interaction | 73 % | 68 % | 72 % | 78 % | 71 % |
| Cell membrane receptors vs.<br>nuclear receptors | 83 % | 80% | 85 % | 88 % | 82 % |
| Receptors of NMD target vs.<br>receptors of non-NMD target | 75 % | 70% | 69 % | 77 % | 70 % |

The second study was to classify cell membrane receptors into either with virus interaction or without one. Recent studies documented that protease functioned as co-receptor to facilitate virus entry and interacted with methyltransferase to modulate the activation of a receptor [25,26]. It was reported that cell membrane receptor bound by a growth factor increased transcript abundances of specific TFs and proteases by the intracellular signaling cascades [27,28]. In addition to data from GUIdEStaR, we included the sign functions of the log base2 FCs of diverse TFs, growth factors, proteases, and methyltransferases. Interestingly, the top attributes that contributed the most to the classification were binary data based upon the sign functions of log base2 FCs in ciliated cells and SINE-mammalian-wide interspersed repeats (MIR). The average accuracy of the binary classifiers was 72%. The third study was to classify the cell membrane and nuclear receptors as either that of nonsense mediated mRNA decay (NMD) target or that of non-NMD target. The top 10 attributes included GQ (4:1:1), long term repeat (LTR) family, Tc1-Mariner-like elements (DNA-TcMar), LINE-L2, DNA-hAT-Blackjack, and m6A modification target sites of METTL14 and YTHDF1, where the GQ format x:y:z was interpreted as “x represents the number of guanine tracts, y represents the number of locations at which a G4 could form, and z represents the number of G4 that could form simultaneously in the sequence” [1]. The average accuracy was 72%. GUIdEStaR is freely available to access the user-friendly web-based database at www.guidestar.kr and to download the entire database and the example Java codes at https://sourceforge.net/projects/guidestar.

## Materials and Methods

### Database Integration

Ensembl gff file (hg38 version: 101) was used to determine the transcript location. A transcript ID (Ensembl ENST) file consists of 845 attributes in the case of mRNA, where each of 5’ UTR, 3’ UTR, exon, intron, and biological region comprises 169 presence/ absence attributes in the order mentioned above. For example, the first 169 presence-absence data of GUIdEStaR were from 5’ UTR region of a transcript: GQ provided by Chariker et al., uORF from uORF-Tools, IRES from IRESbase, epigenetic modifications to the histone protein H3 provided by Choi et al., target sites of m6A writers (KIAA1429, METTL3, METTL14, and WTAP), readers (YTHDC1 and YTHDF1), and eraser (FTO) in 5’ UTR, CDS, 3’ UTR, mRNA, lncRNA, sncRNA, tRNA, miRNA, and tumor suppressor miRNA (miRNATS) from Trew database of Met-DB V2.0, small RNAs from DASHR 2.0, and repeats from RepeatMasker. If a transcript is an lncRNA, it contains three regions in the order of exon, intron, and biological region. G-quadruplex, uORF, IRES, small RNA, and repeats have different binary values per region and per transcript, while epigenetic information (Trew database) has a different value per region and per gene. Epigenetic modifications to the histone protein H3 (H3K27me3 and H3K4me3) have the same values in all five regions of a gene. In three example studies, the example Java codes generated datasets that were readily inputted to neural network methods (Supplementary Data: Example_JAVA_codes).

### Implementation of the web-based database

Java Servlet and HTML with Javascript were employed to implement the database querying and retrieval system. At the homepage, www.guidestar.kr, a user is allowed to submit gene symbols to the system by listing genes in text box and clicking “Send” button. Then, the Servlet program searches user’s gene symbols against the integrated database, retrieves the presence-absence data per region, compresses result files, and sends the compressed zip file to the user if he/ she chooses the option. And it generates an HTML page that displays the retrieved data and a “Download” button that enables the user to save the results in his/her local computer.

**Fig1.**
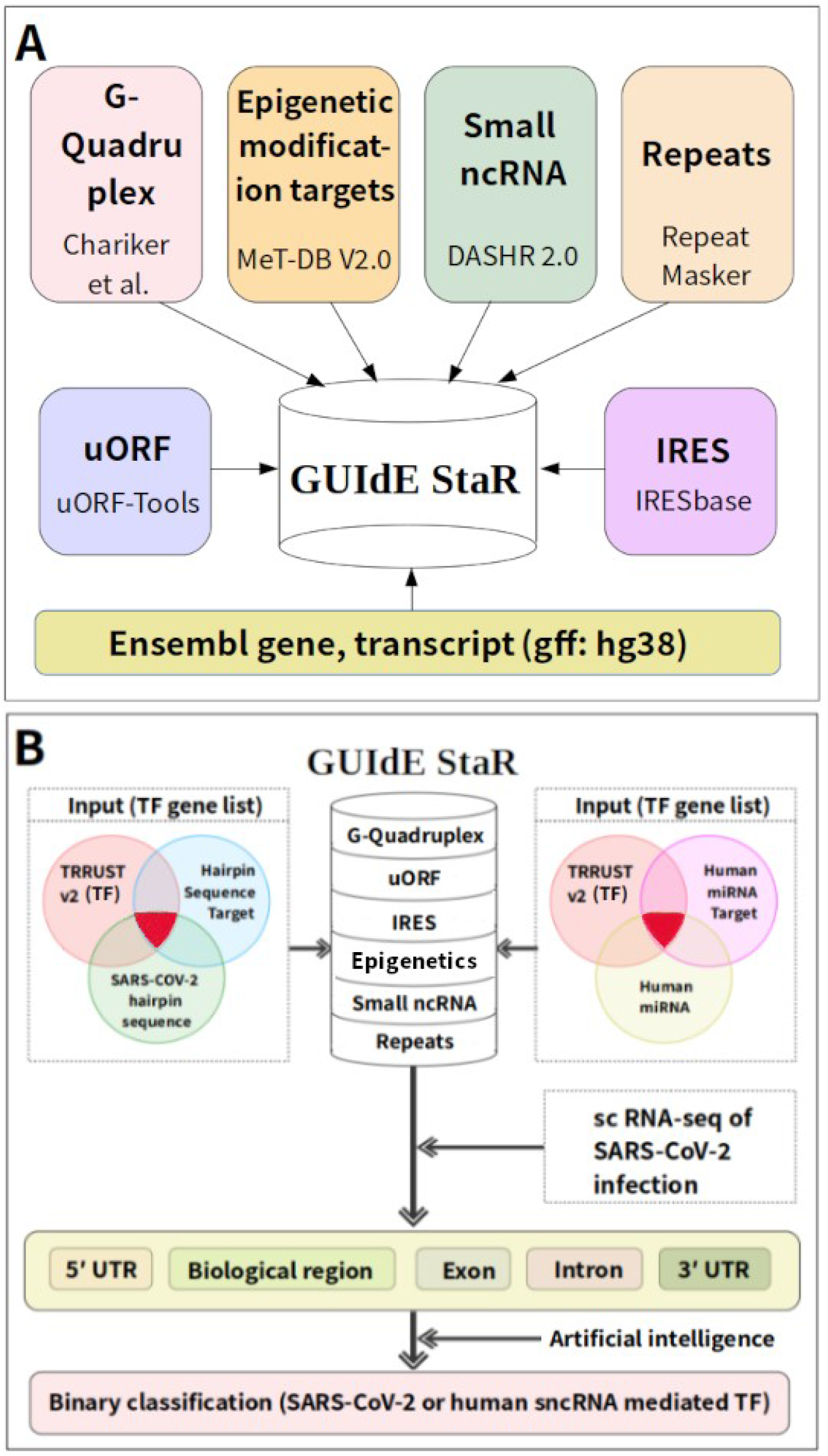
(A) Data integration of GUIdEStaR. (B) Workflow of the small RNA study

### Database usage demonstration

In all three studies, we retrieved the presence/ absence data of 845 attributes from each of all mRNA transcripts belonging to a gene, splitted them into 5’ UTR, 3’ UTR, exon, intron, and biological region, set an attribute to have the value-“true” if the attribute was present in at least one transcript, and included additional scRNA-seq data of SARS-CoV-2 infection, log base2 FC information, in eight different cell types (Supplementary Data: neural_network_inputs) [24]. We built binary classifiers using Weka’s meta classifier, MultiClassClassifierUpdateable with default values (method: against-all, random number seed: 1, base classifier: Stochastic Gradient Descent (SGD), loss function: hinge loss (SVM), learning rate: 0.01, lambda: 1.0E-4, epochs: 500, epsilon: 0.001, normalization) [30]. Resampling preprocessing step was applied in prior to running each classifier due to the small sample number. We assessed the contribution of an attribute to the classification accuracy, using CorrelationAttributeEval with default values for all parameters [30]. For receptor studies, additional 1048 binary values were generated by the sign function that evaluated whether x and y had the same sign where x and y were the log base2 FCs of a receptor gene (subject) and of an attribute-gene family (factor) in a dataset. An attribute gene belonged to one of the following families: growth factor families (bone morphogenetic protein, ciliary neurotrophic factor, LIF, interleukin, epidermal growth factor, erythropoietin, fibroblast growth factor, GDNF family, IGF like family, macrophage, nerve growth factor, neurotrophin, platelet derived growth factor, T cell, megakaryocyte, transforming, and tumor necrosis factor), TF families (AP-2, ARID/BRIGHT, AT hook, BED ZF, Brinker, C2H2 ZF, and other 67 families), proteases (peptidase, transamidase, retropepsin, separase, elastase, glutamyltransferase, and other 51 families), 8 different types of histone methyltransferases (ASH1L, DOT1L, EHMT1, MLL, NSD1, PRDM2, SET, and SETMAR, and other 29 genes), 4 different types of DNA methyltransferases (DNMT1, TPMTP3, DPY30, MGMT, and other 13 genes), 3 different types of RNA methyltransferases (METTL14, NSUN2, TRDMT1, VIRMA, and 87 other genes), or other 10 different types of methylation enzymes (ICMT, SHMT1P1, PCMT1, CARM1P1, EHMT1, CMTR1, AP000812, ARMT1, AS3MT, AC006022, and other 103 genes) (Table S2, JAVA_code_examples: input_files_to_codes)

### Small RNA study

Small RNAs originated from SARS-CoV-2 and from human were selected based on the study of Demirci et al. [31]. TF target genes mediated by SARS-CoV-2 encoded small RNAs were selected based on the same study, while those mediated by human miRNA were based on miRDB and TRRUST v2 (Supplementary Data: neural_network_inputs: small_RNA) [32,33].

### Cell membrane receptor with virus interaction study

Cell membrane receptors that were known to bind to growth factors were selected based upon Wikipedia (https://en.wikipedia.org/wiki/Cell_surface_receptor). Among the receptors, those interacting with different types of viruses were selected based upon the database, Viral Receptor (Supplementary Data: neural_network_inputs: receptor_virus_non_virus) [34].

### Cell membrane and nuclear receptors of NMD target vs. non-target

Nuclear receptors were selected based on wikipedia (https://en.wikipedia.org/wiki/Nuclear_receptor). Genes containing NMD-target transcripts were selected based upon gene annotation of the gff file from Ensembl Homo sapiens 101 (Supplementary Data: neural_network_inputs: receptor_NMD_non_NMD).

## Results and Discussion

The accuracies of binary classifiers estimated with cross-validation (folds=10) are shown in Table 1 (Table S1, Supplementary Data: neural_network_outputs). The top 10 attributes in the small RNA study included epigenetic modification to Histone H3, target sites of m6A writers (WTAP target site overlapping with lncRNA, WTAP with mRNA, METTL14 with miRNATS), repeats (SINE families and LINE families), and log base2 FC data (basal cells, tuft cells, and ciliated cells of infected/bystander cell data) (Supplementary data).

It was interesting to find that the top 10 attributes in the cell membrane receptor with virus interaction study were gene expression comparisons only from ciliated cells: sign functions of log base2 FCs of the following families-protease family (nicalin), transcription factor family (NFX), and growth factor families (erythropoietin, epidermal growth factor, ciliary neurotrophic factor, fibroblast growth factor, GDNF family, neurotrophin, T cell, platelet derived growth factor). It was consistent with the scRNA-seq study that ciliated cells were mainly the targets of SARS-COV-2 infection [24]. Top 10 attributes in the receptors of NMD target study included repeats (LTR, DNA-TcMar, LTR-endogenous retrovirus1 (LTR-ERV1), LINE-L2, DNA-hAT-Blackjack, LTR-ERVL-MaLR), RNA, and target sites of m6A writer and reader (METTL14 target site overlapping with mRNA, METTL14 with 3’ UTR, YTHDF1 with mRNA).

Although omics data availability rapidly increases, the bottleneck is the lack of bioinformatics software and databases that provide users with a convenient means to extract meaningful information that conveys valuable knowledge. GUIdEStaR provides an easy access to key elements of a transcript/gene with a single inquiry. Due to the highly dynamic nature of epigenetic modification, neural network methods such as association rule may reveal useful insights for studying requirements interdependencies of the elements in a transcript and a link between particular features and a specific condition of cellular microenvironment.

## Conclusions

Despite the importance of G-quadruplex, uORF, IRES, Epigenetics, Small RNA, and Repeats, no single resource is available for accessing these information with transcript/gene symbol. GUIdEStaR is the first database that integrates the information into a single data bank, and enables users to retrieve data from database with an easy-to-use query interface. Artificial intelligence methods are nowadays widely used in various fields of biological sciences. GUIdEStaR generates binary datasets, inputs to neural network methods, to promote artificial intelligence driven biological investigations. Here, we also provide a glimpse of the database usage in conjunction with neural network methods; we demonstrate the feasibility of using these six elements as discriminative features to identify human TFs and cell membrane receptors that interact with sequences of viral origin in addition to the identification of receptors of NMD target. We plan to integrate sequence composition and epigenetic modification at single nucleotide resolution into the current GUIdEStaR to provide a useful insight for studying the molecular mechanisms underlying the requirements interdependencies in gene regulation.

## Data Availability

GUIdEStaR is freely available as web-based database at www.guidestar.kr. Additionally, users can download the entire database and the example Java codes from the ‘Download’ page, or at https://sourceforge.net/projects/guidestar/files/.

## Conflicts of Interest

The authors declare no competing interests.

## Funding Statement

This work was supported by grants from National Research Foundation of Korea (NRF-2020R1A6A3A01099714).

## Supplementary Materials

- Table S1. Evaluation of the binary classifiers: https://sourceforge.net/projects/guidestar/files/supplementary_table.zip/download
- Table S2. Families of TF, histone methyltransferase, DNA methyltransferase, RNA methyltransferase, and other methylation enzyme used in receptor study datasets: https://sourceforge.net/projects/guidestar/files/supplementary_table.zip/download
- Example_Java_codes: customized GUIdEStaR database, Java codes and input_files_to_codes: https://sourceforge.net/projects/guidestar/files/Example_JAVA_codes.zip/download
- Supplementary_data: neural_network_inputs and neural_network_outputs (small_RNA, receptor_virus_non_virus, receptor_cell_memb_nuclear, and receptor_NMD_non_NMD): https://sourceforge.net/projects/guidestar/files/supplementary_data.zip/download

## Notes

http://www.guidestar.kr

### Competing Interest Statement

The authors have declared no competing interest.

### Summary of Updates

Web based database with user friendly interface is now available at guidestar.kr

