## supplementary tables for "GUIdEStaR (G-quadruplex, uORF, IRES, Epigenetics, Small RNA, Repeats), the integrated metadatabase in conjunction with neural network methods": TableS1.docx

**Table S1. Evaluation of the binary classifiers**

Small RNA targets (virus origin vs human origin)

(cross validation folds = 10, N= 56)

|  | 5'UTR | Biological region | Exon | Intron | 3'UTR |
| --- | --- | --- | --- | --- | --- |
| virus_origin | 4/7 | 4/7 | 4/7 | 4/7 | 4/7 |
| human_origin | 45/49 | 45/49 | 48/49 | 47/49 | 46/49 |
| Accuracy | 88 % | 88 % | 93 % | 91 % | 89 % |
| Kappa statistic | 0.462 | 0.462 | 0.628 | 0.565 | 0.510 |
| Root mean squared error | 0.354 | 0.354 | 0.267 | 0.299 | 0.327 |

Receptors with virus interaction vs receptors without virus interaction

(cross validation folds = 10, N= 109)

|  | 5'UTR | Biological region | Exon | Intron | 3'UTR |
| --- | --- | --- | --- | --- | --- |
| Virus (correct predictions) | 48/60 | 47/60 | 44/60 | 50/60 | 41/60 |
| Non-virus  (correct predictions) | 32/49 | 27/49 | 35/49 | 35/49 | 36/49 |
| Accuracy | 73 % | 68 % | 72 % | 78 % | 71 % |
| Kappa statistic | 0.457 | 0.340 | 0.446 | 0.552 | 0.413 |
| Root mean squared error | 0.516 | 0.567 | 0.525 | 0.469 | 0.542 |

Cell membrane receptors vs nuclear receptors

(cross validation folds = 10, N= 142)

|  | 5'UTR | Biological region | Exon | Intron | 3'UTR |
| --- | --- | --- | --- | --- | --- |
| Cell membrane receptor | 96/109 | 99/109 | 95/109 | 99/109 | 96/109 |
| Nuclear receptor | 22/33 | 14/33 | 26/33 | 26/33 | 21/33 |
| Accuracy | 83 % | 80% | 85 % | 88 % | 82 % |
| Kappa statistic | 0.536 | 0.367 | 0.614 | 0.675 | 0.512 |
| Root mean squared error | 0.411 | 0.452 | 0.385 | 0.346 | 0.420 |

Receptors of NMD target vs receptors of non-NMD target

(cross validation folds = 10, N= 142)

|  | 5'UTR | Biological region | Exon | Intron | 3'UTR |
| --- | --- | --- | --- | --- | --- |
| Receptors of NMD target | 46/63 | 39/63 | 39/63 | 49/63 | 38/63 |
| Receptors of non-NMD target | 60/79 | 61/79 | 59/79 | 61/79 | 61/79 |
| Accuracy | 75 % | 70% | 69 % | 77 % | 70 % |
| Kappa statistic | 0.488 | 0.395 | 0.368 | 0.546 | 0.380 |
| Root mean squared error | 0.504 | 0.544 | 0.557 | 0.475 | 0.550 |

Note: Filter, resample was applied in preprocess step once due to the small sample number in prior to application of the binary classifiers.
