## supplementary tables for "GUIdEStaR (G-quadruplex, uORF, IRES, Epigenetics, Small RNA, Repeats), the integrated metadatabase in conjunction with neural network methods": TableS2.docx

#### **Table S2. Families of TF, histone methyltransferase, DNA methyltransferase, RNA methyltransferase, and other methylation enzyme used in receptor study datasets**

**TF families:** AP-2, ARID/BRIGHT, AT_hook, BED_ZF, Brinker, C2H2_ZF, C2H2_ZF_AT_hook, C2H2_ZF_BED_ZF, C2H2_ZF_Homeodomain, C2H2_ZF_Myb/SANT, CBF/NF-Y, CCCH_ZF, CENPB, CG-1, CSD, CSL, CUT_Homeodomain, CxxC, CxxC_AT_hook, DM, E2F, EBF1, Ets, Ets_AT_hook, FLYWCH, FYVE-type_ZF, Forkhead, GATA, GCM, GTF2I-like, Grainyhead, HMG/Sox, HMG/Sox_bHLH, HSF, Homeodomain, Homeodomain_POU, Homeodomain_Paired_box, IRF, LYAR-type_C2H2_ZF, MADF, MADS_box, MBD, MBD_AT_hook, MBD_CxxC_ZF, MYM-type_ZF, MYND-type_ZF, Myb/SANT, Myb/SANT_GATA, NFX, NOA36-type_ZF, Ndt80/PhoG, Nuclear_receptor, Paired_box, Pipsqueak, Prospero, RFX, Rel, Runt, SAND, SART-1, SBP, SMAD, STAT, T-box, TBP, TCR/CxC, TEA, THAP_finger, Unknown, ZZ-type_ZF, bHLH, bZIP, mTERF, p53

**Histone methyltransferase types:** histone methyltransferase type1 (ASH1L), histone methyltransferase type2 (DOT1L), histone methyltransferase type3 (EHMT1, EHMT2, EZH1, EZH2), histone methyltransferase type4 (MLL, MLL2, MLL3, MLL4, MLL5), histone methyltransferase type5 (NSD1), histone methyltransferase type6 (PRDM2), histone methyltransferase type7 (SET, SETBP1, SETD1A, SETD1B, SETD2, SETD3, SETD4, SETD5, SETD6, SETD7, SETD8, SETD9, SETDB1, SETDB2), histone methyltransferase type8 (SETMAR, SMYD1, SMYD2, SMYD3, SMYD4, SMYD5, SUV39H1, SUV39H2, SUV420H1, SUV420H2)

**DNA methyltransferase types:** DNA methyltransferase type1 (DNMT1, DNMT3, DMAP1, DNMT3A, FP565260, DNMT3AP1, DNMT3B, DNMT3L, N6AMT1), DNA methyltransferase type2 (TPMTP3, TPMTP4, TPMT, TPMTP2), DNA methyltransferase type3 (DPY30, GAMT, AP003499), DNA methyltransferase type4 (MGMT)

**RNA methyltransferase types:** RNA methyltransferase type1 (METTL1, METTL11B, METTL14, METTL15, METTL15P1, METTL15P3, METTL16, METTL17, METTL18, METTL21A, METTL21AP1, METTL21C, METTL21EP, METTL22, METTL23, METTL24, METTL25, METTL26, METTL27, METTL2A, METTL2B, METTL3, METTL4, METTL5, METTL6, METTL7A, METTL7AP1, METTL7B, METTL8, METTL8P1, METTL9, NSUN2, NSUN3, NSUN4, NSUN5, NSUN6, NSUN7), RNA methyltransferase type2 (TRDMT1, ALKBH8, TRMT1, TRMT10A, TRMT10B, TRMT10BP1, TRMT10C, TRMT11, TRMT112, TRMT112P1, TRMT112P2, TRMT112P3, TRMT112P4, TRMT112P5, TRMT112P6, TRMT112P7, TRMT12, TRMT13, TRMT1L, TRMT2A, TRMT2B, TRMT44, TRMT5, TRMT6, TRMT61A, TRMT61B, TRMT9B), RNA methyltransferase type3 (AC112778, VIRMA, CMTR1, HENMT1, RNMTL1P1, AP004247, BCDIN3D, RAMAC, MRM3, MRM1, FTSJ3, RNMT, TRMU, DIMT1, RAMACL, MRM2, RNMTL1P2, BUD23, BMT2, TRMO, FTSJ1, RNMTL1P1, RNMTL1P2, EMG1, GAMTP2, CMTR2, SPOUT1)

**Other methylation enzymes:**

Cysteine: ICMT, MTR, MTRR, BHMT2, BHMT, GNMT, AC022709

Serine: SHMT1P1, SHMT1

Aspartate: PCMT1, PCMTD1, PCMTD1P1, PCMTD1P2, PCMTD1P3, PCMTD1P7, PCMTD2, PCMTD2, PCMTD1P7, PCMTD1P2, PCMTD1P3, PCMTD1, PCMTD1P1, PCMT1

Arginine: CARM1P1, PRMT1, PRMT1P1, PRMT2, PRMT3, PRMT5, PRMT5P1, PRMT6, PRMT7, PRMT8, PRMT9

Lysine: EHMT1, NTMT1, SETDB1, SETDB2, ASH1L, ASH2L, EEF1AKMT1, EEF1AKMT2, EEF1AKMT3, EEF1AKMT4, EEF1AKNMT, RBBP5, MLLT10, KMT2A, CSKMT, KMT5B, KMT5B, SETD1B, KMT5A, ETFBKMT, KMT2D, SETDB2, AL356585, VCPKMT, SETD1A, EEF2KMT, SETD6, ANTKMT, DOT1L, KMT2B, KMT5C, CAMKMT, AC006453, SETD2, SETD7, ATPSCKMT, EHMT2, KMT2E, KMT2C, ASH2L, EHMT1, KMT5B, SETD1B, ETFBKMT, KMT2D

Leucine: AP000812, LRTOMT, LCMT2, LCMT1, AC133552

Cap: CMTR1, CMTR2

Acidic1: ARMT1

Acidic2: AS3MT, COMTD1, NNMT, COQ5, PEMT, PNMT, HNMT, COMT, AMT, COQ3, INMT, AC006022, CARNMT1, ASMTL

Indolethylamine: AC006022, AC006453, AC006530, AC021220, AC022709, AC024558, AC068305, AC073476, AC112778, AC114781, AC133552, AC239859, AC245595
